# Rapid, reversible RNA/DNA precipitation using rare earth ion coordination

**DOI:** 10.64898/2026.10.12.758480

**Authors:** N. Patel, P. G. Stockley

## Abstract

Efficient recovery and removal of nucleic acids remain major challenges in molecular biology, diagnostics, and viral-vector manufacturing. Conventional precipitation methods often require organic solvents, low temperatures, and extended centrifugation, while nuclease-based contaminant removal can be incomplete and/or is incompatible with downstream workflows. Here, we evaluate NADA, a trivalent rare earth-based reagent, for rapid, solvent-free precipitation of RNA and DNA, and ALGO-mediated recovery of the full range of properties of unmodified RNA. Using hepatitis B virus pre-genomic RNA, lambda bacteriophage DNA, and the HBV nucleocapsid as model systems, we show that NADA rapidly sequesters both ssRNA and dsDNA from solution, reducing Midori Green nucleic acid fluorescence and UV absorption to baseline levels. The NADA complex prevents HBV core protein binding to the HBV pgRNA, and therefore triggering of nucleocapsid formation, even when added simultaneously with core protein, suggesting that lanthanide ion coordination, out-competes the cognate Cp for viral RNA-binding interactions. Recovery with ALGO restores pgRNA integrity and functionality, enabling subsequent assembly of the expected *T*=4/T=3 nucleocapsids. In contrast, pre-assembled nucleocapsids resist NADA-ALGO treatment, demonstrating coat protein capsid-mediated protection of encapsidated pgRNA. Similar outcomes for lambda dsDNA establish NADA/ALGO as a rapid, reversible, and selective alternative platform for nucleic acid precipitation, with potential applications in viral particle purification, contaminant removal, and RNA handling workflows.

## Introduction

Efficient handling and recovery of nucleic acids, especially those with RNA backbones, remains a significant bottleneck in molecular biology and clinical diagnostics^1,2^. Commonly used techniques for nucleic acid precipitation rely on “salting out” the polyanionic backbone of the molecule using monovalent cations (Na^+^, NH_4_^+^), in combination with organic solvents ^3,4^. These methods require the use of sub-zero temperatures and lengthy centrifugation steps. They are prone to both low recovery of dilute samples ^1–4^, and carry-over of organic solvents, which can act as enzymatic inhibitors for downstream applications^4^.

Gene therapies often require the production of pure Virus-Like Particles (VLPs) and Adeno-Associated Virus (AAV) vectors. Such therapeutics are typically manufactured from large-scale mammalian, or insect, cell cultures^5^. This process often generates samples with a high proportion of host cell contaminants^6^. Non-encapsidated host cell DNAs and RNAs can also present significant hurdles as they can trigger innate immune responses in patients, leading to inflammation and reduced efficacy^6^. Current removal methods for such contaminants often rely on the use of nucleases, such as Benzonase^7^ to digest contaminant nucleic acids. However, such digestion can be inconsistent, requires to be used in specific buffer conditions, and leaves behind small fragments that are difficult to remove during chromatography^6–8^. There is a need for purification technologies that can selectively and rapidly sequester free nucleic acids.

Here we used NADA^9,10^, a reagent available from Entropix Ltd., that contains a trivalent rare earth ion. This reagent uses Lewis acid coordination chemistry to achieve rapid, solvent-free RNA or DNA precipitation. Its accompanying ALGO solution then facilitates unmodified nucleic acid recovery. Trivalent rare earth ions exhibit a coordination affinity for phosphates which exceeds the electrostatic interactions of more traditionally used cations in nucleic acid extractions^11^. As previously described ^12,13^, rare earth ions can form inner-sphere coordination complexes, that displace the hydration shell of the nucleic acid and induce a “charge inversion”, resulting in a solid, positively-charged RNA pellet^12–16^.

To explore the binding and diagnostic utility of this technology, we used the Hepatitis B Virus (HBV) nucleocapsid (NC) as a biological model (Fig. 1). The assembly of HBV into *T*=4 NCs is a well-characterised process governed by the recognition of distributed single-stranded RNA packaging signals by the arginine-rich C-terminal domains (CTDs) of the viral Core Proteins (Cps) ^17–19^. HBV NCs are known to shield their internal RNA from chemical or enzymatic access^20–22^.

**Figure 1:**
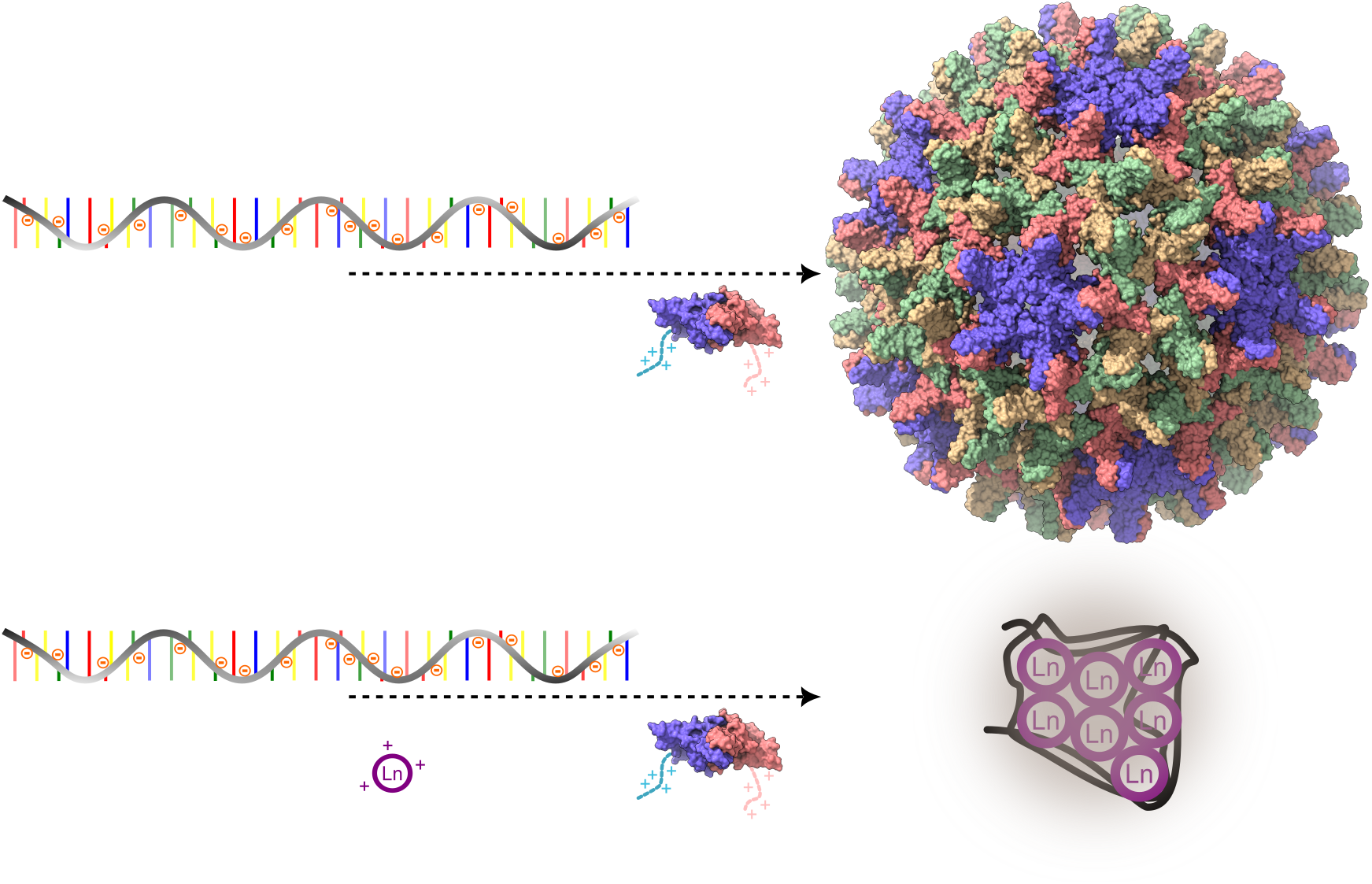
An HBV NC assembly system. *Top*, The negatively-charged phosphodiester backbone (*left, black*) of the pgRNA interacts with the positively-charged RNA binding domain (an arginine-rich domain (ARD)) of the HBV core protein (Cp) (*red/blue/orange/green*) forming a *T*=4 nucleocapsid (*right*) comprised of 240 Cp identical subunits (*red/blue/orange/green*) ^17, 22^. *Bottom*, upon introduction of 3 µL of the rare earth ion-based 50x NADA (exemplified by Ln^3+^ (*purple*), prior to the addition of HBV Cp, the rare earth ions bind to the pgRNA and cause it to precipitate, preventing formation of the HBV nucleocapsid.

## Methods

### Nucleic Acid Binding and Displacement Assays

HBV pgRNA was produced via *in vitro* transcription using the HiScribe T7 High Yield RNA Synthesis Kit (New England Biolabs) according to the manufacturer’s instructions and purified as previously described^17,22^. 1 µL Midori Green Direct (*Nippon Genetics*), a non-intercalating dye that associates with the polyanionic backbone of nucleic acids was added to 1 µg of either the Entropix supplied λ DNA (dsDNA) or HBV pgRNA (ssRNA) in a buffer containing 50 mM HEPES (pH 7.4) and 250 mM NaCl, at a final volume of 50 µL.

According to the manufacturer’s instructions, 50x NADA was added to this solution in 1 µL additions up to 4 µL, and fluorescence emission spectra (ex/em 490/520 nm) were recorded at room temperature using a FLUOstar Omega plate-reader (BMG Labtech).

### HBV Nucleocapsid Assembly Assays

HBV core protein (Cp) dimers, purified as previously described ^17,22,23^, were used to assemble nucleocapsid-like particles (NCs) around the HBV pgRNA. The latter was produced as previously described ^17^. 1 – 4 µL of 8.5 µM Cp dimer were then added to 18 nM of pgRNA in 50 µL total volume. This meant dimers were added at a molar ratio of 120 (1 µL), 240 (2 µL), 360 (3 µL) and 480 (4 µL) Cp: 1 pgRNA, and then incubated at room temperature for 30 min, ensuring the assembly of complete NCs.

Assembly was assessed under five conditions: 1) as described above in the absence of NADA; 2) pgRNA was pre-incubated with 3 µL 50x NADA before Cp addition as above; 3) 4 µL Cp and 3 µL 50x NADA were added to pgRNA concurrently using separate pipettes to prevent Cp precipitation; 4) pgRNA was recovered according to the manufacturer’s protocol using the supplied cellulose acetate filters and ALGO solution. HBV pgRNA was cleaned prior to use using Monarch RNA purification columns, according to the manufacturer’s protocol (*New England Biolabs)*, and subjected to the above assembly assay; *5)* HBV NCs were assembled, as described above, before the addition of 3 µL 50x NADA.

### Characterisation of NC Assembly

Following treatment and incubation with 50x NADA, samples were centrifuged at 16,000 x g for 1 min to pellet any insoluble aggregates. The A_260/280_ ratio of the resulting supernatant was measured using a NanoDrop™ One (*ThermoFisher Scientific*), and a value of ∼1.55 was used to represent the characteristic protein-to-nucleic acid stoichiometry of a *T*=4 capsid^17^. Absorbance values were corrected for light-scattering using the A_310_ and A_340_ values, as described^24^.

5 µL of each sample was applied to glow-discharged carbon-coated copper grids and negatively-stained with 2% (w/v) uranyl acetate. Grids were visualized using a Talos L120C transmission electron microscope (*ThermoFisher Scientific*) operating at 120 kV, at 56,000x magnification (Fig . 3).

Protein and RNA integrity within these reactions were confirmed via 15% (w/v) SDS-PAGE and 1% (w/v) denaturing formaldehyde agarose gel electrophoresis, respectively (Figs. 4A & B).

## Results

### NADA induces rapid precipitation of nucleic acids

#### a) ssRNA

A solution containing HBV pgRNA exhibited robust fluorescence when bound to Midori Green (2401 RFUs, Table 1). Upon addition of NADA, followed by centrifugation to remove precipitants, fluorescence from the solution was quenched to baseline levels (Table 1), indicating that the rare earth-based reagent effectively precipitates the nucleic acid from solution (Fig. 2). These titrations confirmed that 3 µL of 50x NADA is sufficient to achieve complete phosphodiester backbone saturation of 1 µg of the ss pgRNA (Fig. 2, Table 1).

**Table 1:** NADA effects on HBV pgRNA and λ dsDNA. *Left to Right*: The sample tested, the treatment of each sample, and the resulting fluorescence (520 nm) and absorbance (260 nm) are listed. Lastly, the outcome of each experiment based on these observations and the nsEM is noted.

| Sample | NADA- Treatment | Fluorescence (RFU) | Absorbance (AU) ($A_{260}$) | Presumed Outcome |
| --- | --- | --- | --- | --- |
| $\lambda$ DNA (dsDNA) | None | 6205 ± 103 | 3.7 | |
| | + 3 μL 50x NALDO | 21 ± 1 | 0.3 | Solution fluorescence quenched |
| pgRNA (ssRNA) | None | 2401 ± 28 | 3.1 | |
| | + 1 μL 50x NADA | 1241 ± 11 | 1.2 | Fluorescence partially quenched |
| | + 3 μL 50x NADA | 150 ± 13 | 0.4 | Fluorescence quenched |
| Pre-assembled NCs | + 3 μL 50x NADA | 3215 ± 42 | 1.3 | Slight fluorescence increase alongside NCP assembly |

**Figure 2:**
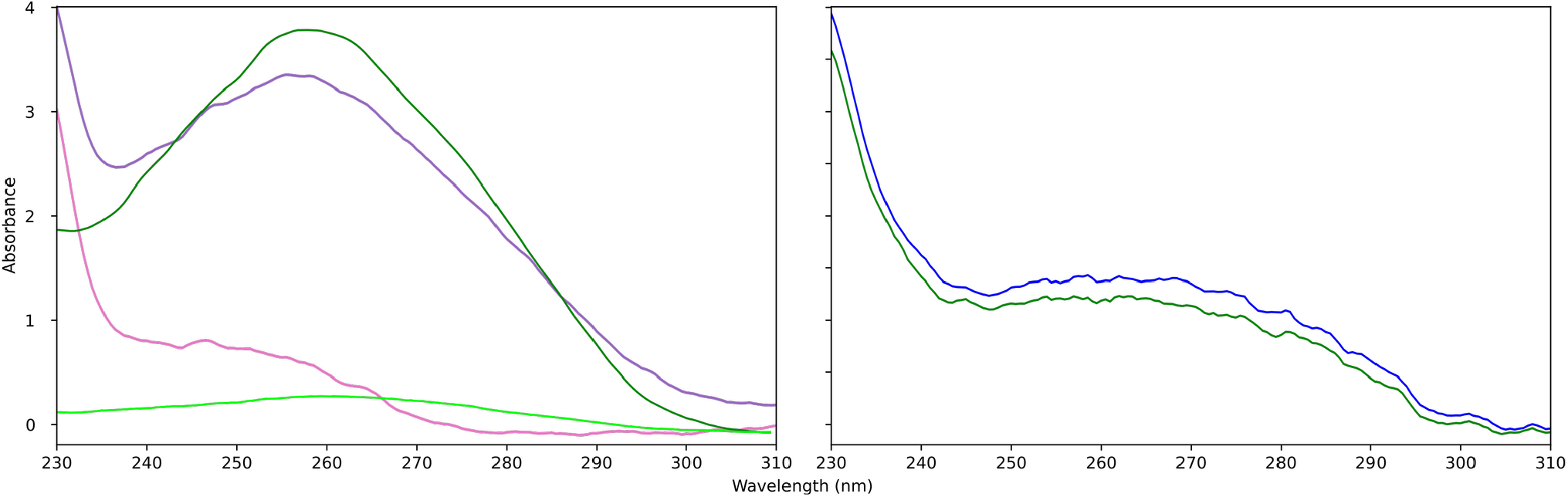
Spectroscopic Assays of NADA/ALGO precipitations. *Left-hand side:* UV spectra of 1 µg pgRNA and 1 µg λ dsDNA (*purple and dark green, respectively*) in the absence and presence (*pink and light green, respectively*) of 3 µL 50x NADO. *Right-hand side*, UV spectra of HBV NCs pre-assembled around 1 µg pgRNA in the absence *(cyan)* or presence *(blue)* of 3 µL 50x NADO.

#### b) dsDNA

Identical titrations were carried out with λ DNA (*Entropix Ltd*.). Fluorescence was observed on binding of the Midori Green to the DNA. This fluorescence was quenched to baseline levels on exposure to the NADA reagent, consistent with complete phosphodiester backbone saturation (Table 1). Absorbance at 260 nm also followed the same trend (Table 1), confirming this interpretation.

### Rare Earth Ion Coordination Inhibits Viral Assembly and Sequesters RNA

Addition of HBV Cp to pgRNA pre-incubated with NADA fails to result in in vitro HBV NC assembly as seen in Table 2, and confirmed by negative stain EM (Fig. 3B). The fluorescence of the pgRNA solution on addition of Midori Green remains at baseline (<200 RFU) compared to the assembly control (∼3500 RFU; Fig. 4A, Table 2). Furthermore, the added Cp remained soluble despite the presence of the rare-earth reagent (Fig. 4A). nsEM confirmed these observations by the presence and absence of NCs, and absorption analysis (A_(260/280)_ ratios) of centrifuged samples were consistent with no nucleic acid remaining in the supernatant (Table 2). These data are consistent with NADA inducing rapid precipitation of the RNA, making it unavailable for HBV Cp binding.

**Table 2:** Impacts of NADA/ALGO on HBV Assembly and pgRNA Recovery. *Left to Right*: The NCP experiment, and the resulting fluorescence (520 nm), absorbance (260 nm) and A260/280 ratios are listed. Lastly, the conclusions drawn as a result of these observations.

| Assembly Experiment | Fluorescence (RFU) | Absorbance (AU)<br>(A<sub>260</sub>) | A<sub>260/280</sub> Ratio | Presumed Outcome |
| --- | --- | --- | --- | --- |
| NCs Assembled as described<sup>17</sup> | 3182 ± 47 | 1.4 | 1.48 ± 0.16 | Normal T=4 NCs produced |
| Pre-incubation of pgRNA with NADA | 134 ± 10 | 0.2 | 0.76 ± 0.05 | No NC assembly; HBV pgRNA pre-bound to NADA |
| Simultaneous addition of Cp + NADA to pgRNA | 109 ± 9 | 0.3 | 0.7 ± 0.05 | NADA outcompetes Cp for binding the pgRNA |
| pgRNA:ALGO recovery of pgRNA | 2325 ± 41 | 1.3 | 1.49 ± 0.11 | T=4 NC assembled around recovered pgRNA |

**Figure 3:**
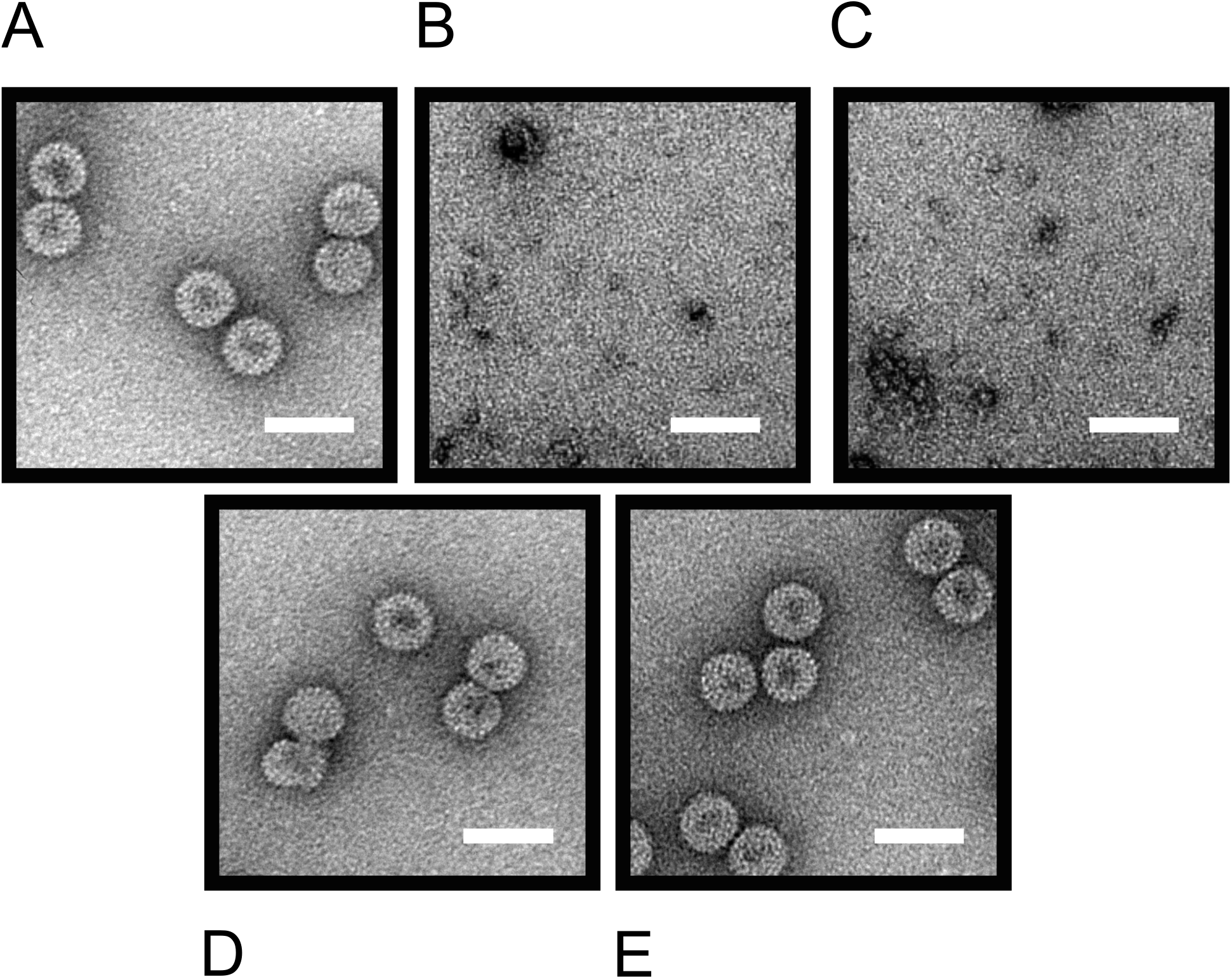
**A)** Wide field nsEM image of the reassembly experiments between HBV Cp and pgRNA. Additional micrographs of the same assembly where **B)** the pgRNA was pre-incubated with 3 µL 50x NADA, **C)** pgRNA was exposed to 3 µL 50x NADA and HBV Cp concurrently, **D)** pgRNA recovered using the ALGO solution was assembled using HBV Cp and **E)** pre-assembled HBV NCs were exposed to 3 µL 50x NADA. All size markers = 50 nm.

In competition assays, when Cp and NADA are added simultaneously from separate solutions, the NADA reagent successfully outcompetes Cp for pgRNA binding, as evidenced by the lack of assembly-dependent or RNA-dependent Midori Green fluorescence (Fig. 3C, Table 2). This conclusion was also confirmed using nsEM.

To determine if the rare earth-induced precipitation of pgRNA was reversible, we used aqueous ALGO solution (*Entropix Ltd*.). Following precipitation and pelleting of the pgRNA:NADA complex, the pellet was resuspended using ALGO (Fig. 4B). Upon release of the rare earth ions, the functionality of the released transcript as an assembly substrate was confirmed using the same Midori Green based assay. Quantitative nsEM analysis and A_260/280_ ratios of the assembly mix (∼1.5, Table 2) confirmed the successful assembly of *T*=4 NCs (Fig. 3D). This confirms that whilst NADA-coordination is high-affinity and effectively outcompetes Cp for binding to pgRNA, it does not covalently modify or degrade the nucleic acid (Fig. 4B). This allows for the rapid concentration and release of viral genomes for subsequent study.

**Figure 4:**
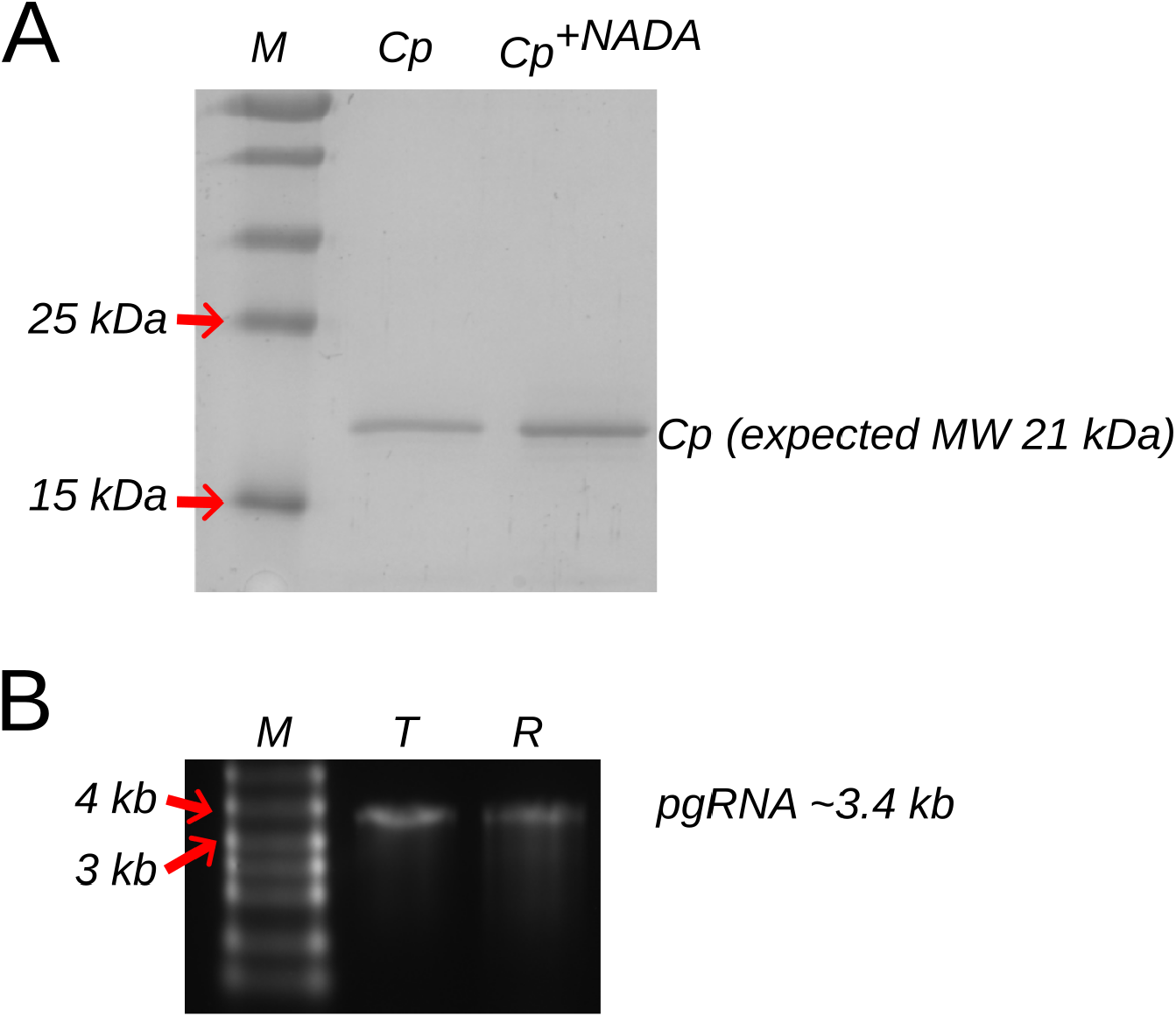
Gel analysis of NADA precipitation. **A)** SDS PAGE gel (15% (*v/v*)) showing the expected dissociated (?) HBV running at the expected molecular weight. M = Protein marker, Cp = dissociated HBV Cp, Cp = dissociated HBV Cp + 3 µL 50x **NADA. (B)** Denaturing 1% (*w/v*) formaldehyde agarose gel, with the pgRNA (from T7 transcription) rescued from the NADA precipitation by subsequent addition of the ALGO solution (R), and (*T*) shown. Size markers are shown (*M*).

### Capsid Encapsidation Provides a Steric Shield Against NADA

In contrast to free RNA, pre-assembled HBV NCs (Fig. 3A) are unaffected by NADA treatment (Fig. 3E). Fluorescence of the NCs remains stable (at ∼3200–3600 RFU; Fig. 2, Table 1), and mirrors results obtained prior to initial NADA precipitation the previous levels of NCs assembled in the absence of NADA. This was confirmed by electron microscopy, nsEM confirming the presence, and persistence, in solution of the expected mixture of *T*=3 and *T*=4 NCs (Methods). These data suggest that the HBV Cp protein shell of the NC acts as a physical barrier, protecting the internal pgRNA from rare earth ion-induced displacement and precipitation.

## Discussion

Our results demonstrate that NADA is a potentially superior free nucleic acid precipitant to commonly used techniques such as those which use divalent ions in combination with alcohols^1,2,3,4^. NADA utilises coordination affinities that out-compete the electrostatic and other interactions of biological protein partners, in this case, the HBV Cp. We used NADA to induce charge inversion and molecular compaction^12,13,16^, preventing the sequence-specific recruitment of viral proteins to their packaging signals^17^.

The technical advantages of the NADA/ALGO system over traditional alcohol-based precipitation or enzymatic digestion are three-fold: 1) Speed and efficiency - quantitative sequestration occurs at room temperature in seconds, offering a significant time reduction in processing; 2) Selectivity - the shielding effect of the NC shell allows differentiation between encapsidated and free pgRNA, but in principal any sequence-specific nucleic acid binding protein. Through simplifying the removal of contaminants, NADA can act as an efficient “cleaning” agent for VLP preparations; 3) Biocompatibility **–** the aqueous ALGO solution ensures that recovered RNA remains functionally competent for further downstream assays, e.g. for verifying the quality and identity of the viral cargoes during vaccine and gene therapy production.

In conclusion, NADA is a versatile tool for both fundamental research and clinical biomanufacturing. Its ability to manipulate the physical state of RNA through coordination chemistry offers a faster, solvent-free alternative for nucleic acid precipitation, which currently commonly relies on alcohol precipitation steps. Furthermore, its selective interaction with RNA excluded from viral particles provides a method for assessing the integrity of, and clean impurities from, complex nucleoprotein assemblies. Its ability to “clean” preparations of VLPs, is a potentially critical step in ensuring the safety and purity of gene therapy and vaccine-based therapeutics.

